# A species difference in glycerol-3-phosphate metabolism reveals metabolic adaptations in hares

**DOI:** 10.1101/2024.04.02.587662

**Authors:** Kateryna Gaertner, Mügen Terzioglu, Craig Michell, Riikka Tapanainen, Jaakko Pohjoismäki, Eric Dufour, Sina Saari

## Abstract

The temperate climate adapted brown hare (*Lepus europaeus*) and the cold-adapted mountain hare (*Lepus timidus*) are evolutionarily closely related and interfertile. Still, their cultured skin fibroblasts show clear differences in the expression of genes related to basic cellular metabolism. To study this further, we utilized targeted metabolomics analysis, metabolite tracing, and high-resolution respirometry, and identified significant differences in metabolic pathways associated with adaptive thermogenesis, including a higher rate of glycerol 3-phosphate (G3P) production in the mountain hare. We therefore investigated mitochondrial heat production and its dependence on G3P in the two hare species. The mountain hare maintained lower mitochondrial temperature and had weaker thermal change following OXPHOS inhibition. Manipulating mitochondrial glycerol 3-phosphate dehydrogenase (GPD2) levels demonstrated its role in mitochondrial thermogenesis and revealed species-specific function in maintaining mitochondrial membrane potential. This study unveils previously undocumented and unexpected species differences in the mitochondrial properties of fibroblasts that could indicate differences in metabolic adaptability. These findings also demonstrate the utility of cell culture models in assessing trait differences between species and their evolutionary significance, contributing to a deeper understanding of metabolic adaptation in animals and underscoring the potential of *in vitro* approaches for eco-physiological studies.

## Introduction

Fibroblasts are one of the most common and essential cell types, shaping the connective tissue that surrounds different organs. Skin fibroblasts are part of the first line of defense against external stressors such as cold (1). They participate in wound healing (2), while providing, maintaining, and remodeling the extracellular matrix (3). Since their central role in tissue maintenance and repair requires plasticity and proliferative capacity, fibroblasts associate with hyperproliferative pathologies such as fibrosis (1) and cancer progression, where their metabolic rewiring contribute to tumor metastases (4).

Although research on wildlife-derived cells remains limited, there is ample evidence to suggest that fibroblasts of different species have evolved distinct features. Studies comparing skin fibroblasts from tropical and temperate bird species have linked higher chemical and oxidative tolerance to longer species lifespans and lower metabolic rates (5,6). Similarly, skin fibroblasts derived from rodent species with different life history exhibit species-specific tolerance to chemical stressors even in animals collected from the same habitat (7,8). These examples demonstrate that cells harbor species-specific traits, plasticity, and metabolic adaptation even when comparing related species living in the same natural habitat.

Understanding metabolic adaptation at the cellular level could provide new insights on how animals can survive in a changing environment (9). In the face of climate change, many ecosystems are rapidly changing, challenging their host species, particularly those that are restricted to the affected habitats. Simultaneously, environmental changes provide favorable conditions for new species to thrive. Finland presents a typical example of a subarctic region, where the continental European brown hare (*Lepus europaeus*) is expanding its range of habitat, thereby competing with the resident mountain hare (*Lepus timidus*) and causing it to retreat northward (10,11). The mountain hare has evolved in the harsh conditions of circumpolar arctic and subarctic habitats, being separated by 3 million years of evolution from the brown hare (12), which in turn has adapted to the temperate regions of open steppe and bushlands of the southern Palearctic (13). It is still unclear whether climate change, human impact, species-specific characteristics (e.g. litter-size differences), or a combination of these factors have enabled the current expansion of the brown hare’s range in northern Europe (11,14).

Given the distinct evolutionary trajectories of the mountain hare and the brown hare, we investigated their cellular metabolic features using skin fibroblasts isolated from a sympatric population of these two hare species in Finland. We identified species-specific differences in the cell physiology of brown and mountain hare fibroblasts (15). Specifically, brown hare fibroblasts showed higher proliferation rates and faster wound closure. At the molecular level these properties were associated with species-specific variation in the duration of the cell cycle phases as well as differences in the electrochemical properties of mitochondria (15). While changes in the cell cycle progression are likely linked to cell proliferation and wound healing capacity, differences in mitochondrial properties like maintenance of membrane potential may be rooted in the metabolic acclimation of these two species of hare.

Mitochondria are metabolic hubs, that act not only as a terminal step for metabolic degradation linked to energy production, but also provide essential biosynthetic intermediates for cell growth and proliferation. In addition, mitochondria act as central signalling organelles, directly and indirectly influencing e.g., cell death, nuclear gene expression via transcription factors and chromatin modifications, as well as metabolic switches from anabolic to catabolic states (16).

Here we reveal differences between the two hare species related to mitochondrial metabolism and bioenergetics. Using targeted metabolomics, we show that cells derived from the mountain hare have significantly higher levels of metabolites known to associate with NADH recycling, energy storage, and adaptive thermogenesis, including glycerol-3-phosphate (G3P) (17,18). Metabolic flux analyses confirm a higher rate of G3P production in mountain hare cells, and high-resolution respirometry confirmed species differences in G3P-driven mitochondrial respiration. In addition, our study unveils previously undocumented and unexpected species differences in mitochondrial thermogenesis and membrane potential associated with G3P energetic metabolism. Altogether, this study reveals unforeseen species differences in fibroblast metabolism, supporting the idea that animal speciation translates into measurable metabolic differences in cell culture.

## Results

Our previous study of brown hare (*Lepus europaeus,* LE) and mountain hare (*Lepus timidus,* LT) fibroblast biological functions identified species-specific differences in cell proliferation, cell cycle and wound healing, and mitochondrial metabolism (15). To better understand the species-specific traits underlying these differences, we performed targeted metabolomics analysis. Out of 147 identified metabolites seventeen were found significantly different between the two species, ten of which were present at higher levels in LE cell lines and seven in LT cell lines (Figure 1A, Table S1). Amongst the seven metabolites present at significantly higher levels in LT cells, were the three components of the creatine kinase system, creatine (Cr), phosphocreatine (PCr), and creatinine. The creatine kinase (CK/PCr) system converts mitochondrial ATP and Cr into PCr which serves as a stock of rapidly deployable energy for the cell (19). Creatinine is a product of the spontaneous degradation of Cr and PCr. Recent studies have identified alternative thermogenic function of the CK/PCr system, namely creatine-dependent thermogenesis in mitochondria of brown adipocytes (reviewed in (20)). However, to our knowledge, such system has not been previously described in skin fibroblasts. Another metabolite linked with the regulation of energy metabolism; glycerol 3-phosphate (G3P) was also present at significantly higher levels in the LT cells. As a substrate for the glycerol-3-phosphate (G3P) shuttle, G3P sits at the crossroad between glycolysis, fatty acid metabolism and oxidative phosphorylation. The G3P shuttle transfers the reducing power from cytosolic NADH to the mitochondrial respiratory chain via G3P dehydrogenase 1 (GPD1, cytosolic) and G3P dehydrogenase 2 (GPD2, mitochondrial) (18). Notably, the use of this G3P shuttle is known to generate heat (18,21). Succinyl- and Malonyl-CoA were present at significantly lower levels in LT cells. Fatty acid synthesis roots in Succinyl-CoA, an intermediate of the tricarboxylic acid cycle (TCA) which provides Coenzyme A for Malonyl-CoA synthesis. Importantly, Malonyl-CoA is the main inhibitor of fatty acid catabolism. Altogether, these observations suggest species-specific differences in creatine metabolism and in OXPHOS activity connected to G3P.

**Figure 1:**
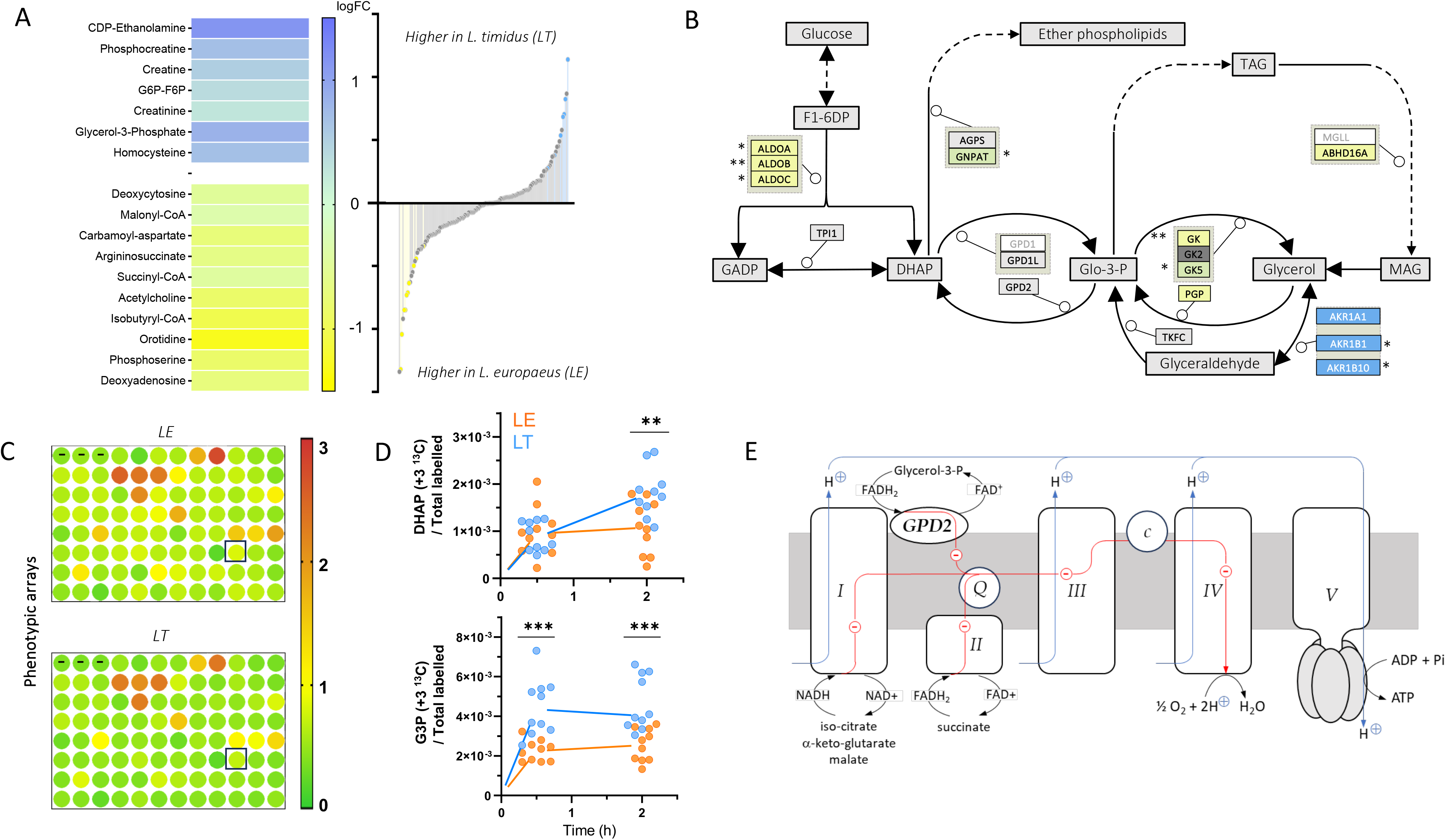
Hare fibroblasts present differences in metabolites related to energy cycling. (A) Seven metabolites are present in significantly higher level in Lepus timidus. On the left side: heatmap clustering analysis of the metabolites present at significantly higher levels (logFC) in L. timidus (LT, in blue) and in L. europaeus (LE, in yellow) fibroblasts. On the right side: distribution of the 147 metabolites identified in LT and LE fibroblast lines. Positive and negative values on the y-axis respectively indicate higher levels in LT and in LE. The significant differences are highlighted (uncorrected Student’s t-test, alpha = 0,05; n = 17/147) and sorted from the most significantly changed at the top and bottom to less significant towards the center. Metabolite levels were corrected based on protein quantitation. n= 6 per fibroblast line (LT 5: n=5; LT 6: n=4), 4 fibroblast lines per species. (B) Transcriptomic analysis suggests differences in G3P metabolism between the two Lepus species. Simplified map of the metabolic pathway centered around G3P. The colour code for highlighting the enzymes is the same as in panel A. * Significant before FDR correction, ** < FDR 0,05. (C) LE fibroblasts present generally higher metabolic rates than LT. NADH buildup in LE and LT fibroblasts. Each dot corresponds to one well of a 96 plate where no substrate (-) or a single metabolite (see Supplementary table 2 for a list), is provided to the cells as the unique source for carbon metabolism. The boxed position corresponds to G3P. The intensity of the signal corresponds to the averaged amount of NADH accumulated after 48h of growth for the given species. N = 4 measures per genotype. (D) LT fibroblasts present higher flux of glucose to G3P. Proportion of ^13^C glucose converted into DHAP (upper panel) and G3P (lower panel) in LT (blue) and LE (yellow) fibroblast cells (calculated as, at each timepoint and for each sample, intensity of the ^13^C-labelled metabolite of interest / sum of the intensities of all measured ^13^C-labelled metabolites). Two cell lines per genotype, 5 measurements per cell line.

These findings prompted us to re-analyze previous transcriptomics results (15) focusing on genes related to G3P- and creatine-metabolism. Some transcripts were detected in one species only (Figure 1B), suggesting an extremely low expression or a strong divergence from the consensus sequences in the other species. In the creatine pathway, the mitochondrial Glycine amidinotransferase (*GATM*) responsible for synthesizing guanidinoacetate, the precursor for creatine, as well as two alkaline phosphatases converting PCr in creatine + P were only found in the LT cells. These species-specific transcriptional differences in the creatine pathway support an increase in the synthesis of creatine and of the cycling between creatine and PCr in the LT cells (Figure S1). Associated to the G3P metabolism, *ARK1A1*, an Aldo-keto reductase capable of converting glycerol into glyceraldehyde was one of the genes found in the LT cells only. *ARK1B1* and *ARK1B10*, the two other enzymes catalyzing this reaction were highly upregulated in LT cells compared to LE. In contrast, *PGP*, which converts G3P to glycerol was found only in the LE samples, and *GK* and *GK5*, which catalyze the opposite reaction were present at higher level in the LE cells compared to LT (Figure 1B). These observations suggest strong differences in the influx and effluxes associated to G3P but fail to explain why G3P itself is present at higher levels in the LT fibroblasts.

We thus compared the ability of the LT and the LE cell lines to grow on various substrates using the Biolog Phenotype MicroArray^TM^ (22). Cells were grown on 96- well plates pre-loaded with a single energy source per well. As the culture medium was mainly devoid of energy sources, cell growth depended on the substrates provided in the wells. Using formazan dye, we periodically measured the cellular NADH levels in each well for 70 hours as a proxy for cell growth and metabolic activity. NADH accumulation was generally weaker in LT cells (Figure 1C), which could be indicative of a lower metabolic efficacy and likely relates to their slower growth rate (15). Nine metabolites provided notably higher NADH levels than the others: D- maltose, a-D-glucose, D-Mannose, Maltotriose, D-Fructose, Inosine, D-Galactose, Uridine and Adenosine (Table S2). This signature shows strong similarities with many human cell lines (22,23). Surprisingly though, G3P substrate was not significantly preferred by the LT cells.

To verify the differences in G3P levels between the two species and assess the connection between G3P levels and glucose metabolism, we performed metabolic tracing using ^13^C glucose, in which all six carbon atoms are replaced by heavy ^13^C. We measured the proportion of heavy carbons transferred from ^13^C glucose to Ribose- 5-P (+5 ^13^C), DHAP (+3 ^13^C), G3P (+3 ^13^C), lactate (+3 ^13^C), citrate, succinate (+2 ^13^C) and phosphoserine (+3 ^13^C) (Figure S2). Upon addition of ^13^C glucose, we observed a rapid surge in G3P labeling followed by an increase in dihydroxyacetone phosphate (DHAP) labeling after 1.5 h in the LT cells (Figure 1D), supporting a glycolytic origin of the G3P and corroborating the metabolomics finding.

G3P is a key intermediate for the recycling of cytosolic NADH to NAD^+^ by the G3P shuttle. This role is intimately linked to the mitochondrial OXPHOS (Figure 1E) *via* the mitochondrial GPD2 enzyme (21). Western blot analysis of GPD2 excluded a differential expression of this protein between the LT and LE fibroblasts (Figure 2A). While GPD2 appeared as a double band, suggesting the existence of a modified or non-processed isoform, each potential isoform was present in similar amounts in both species. In contrast, species-specific GPD2 coding sequence differences were found at position 187 and 241 (Figure S3A). Compared to the asparagine in the mountain hare, the presence of a serine at position 187 in the brown hare GPD2 caused a lowered homology with the consensus motif for FAD dependent oxidoreductases (Figure S3B), possibly indicative of lower catalytic activity.

**Figure 2:**
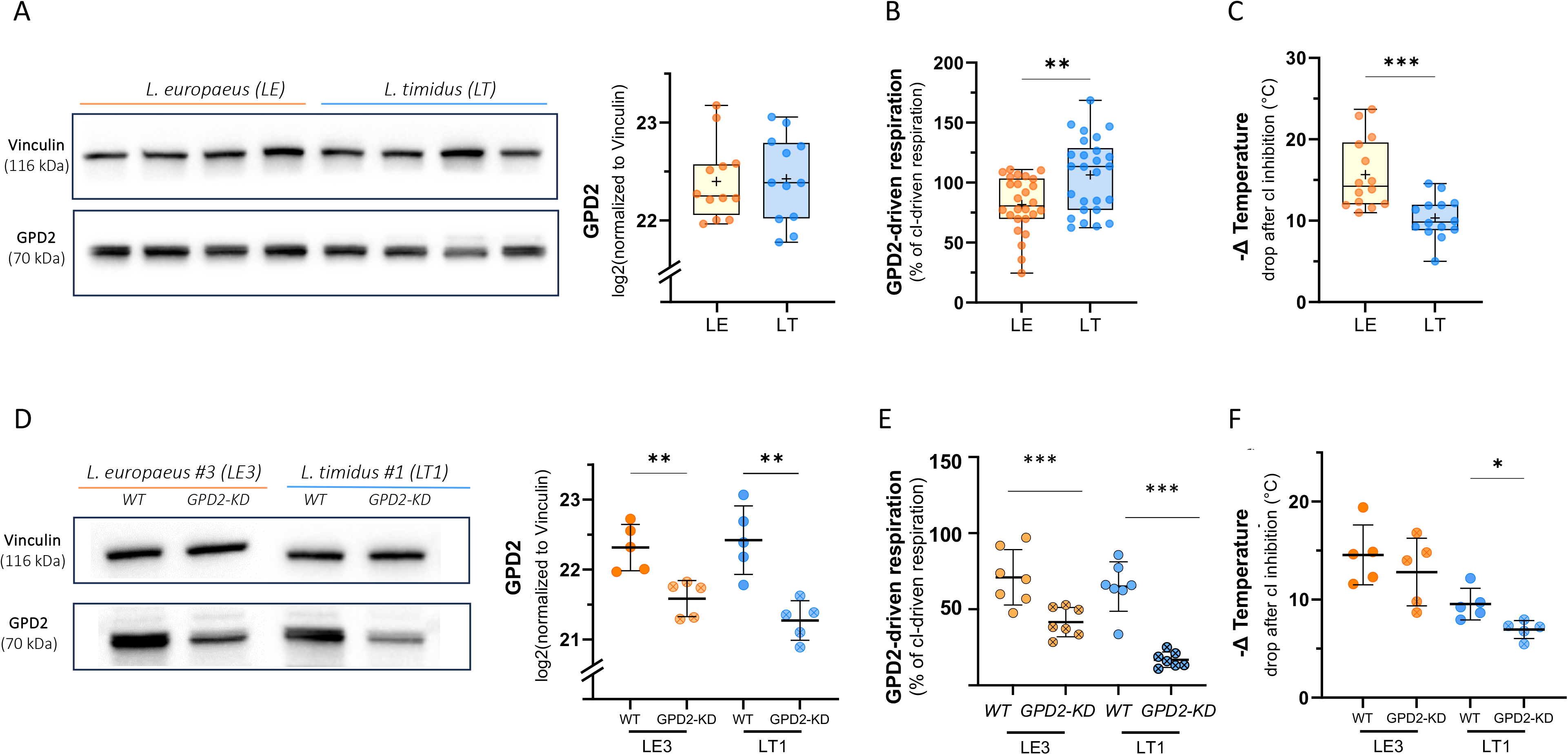
GPD2 activity is higher in LT fibroblasts. (A) Western blot quantitation of GPD2 protein in LE and LT fibroblasts. No significant differences were detected in the protein level of GPD2 between the two species. (B) GPD2-driven respiration is higher in LT fibroblasts. (C) LT fibroblasts have lower decrease in mitochondrial temperature than LE cells after Complex I inhibition with rotenone. (D) Western blot quantitation of GPD2 protein in LE and LT fibroblasts with GPD2 knock-down. Protein level of GPD2 was significantly lower in both LE and LT GPD2-KD cell lines. (E) GPD2-driven respiration in GPD2-KD cell lines is significantly decreased in both LE and LT GPD2- KD cell lines. (F) GPD2 knock-down leads to lower decrease in mitochondrial temperature in both LE and LT cells after Complex I inhibition with rotenone compared to WT cells. The decrease was only found significant in LT cells. *p-value < 0.01, **p- value < 0.001, ***p-value < 0.0001. WT = wild type.

GPD2 converts G3P into DHAP through a reaction coupled to the mitochondrial OXPHOS. This allows the measurement of GPD2 activity by recording the oxygen consumption (OCR) in the presence of exogenous G3P and ADP (Figure 1E). To correct for the imprecision in cell count and potential variations in OXPHOS content per cells, G3P-dependent respiration was normalized to either Complex I- or to Complex IV-driven respiration. G3P-driven respiration was ∼25% higher in the LT permeabilized fibroblasts, whether normalized to Complex I- (Figure 2B) or complex IV-driven (Figure S4A) respiration, indicating an increased capacity to use G3P as a respiratory substrate. This difference was further increased in the absence of normalization (∼38%, Figure S4B). High activity of GPD2 in cultured fibroblasts has historically been linked to their high glycolytic metabolism (24). However, this hypothesis is inconsistent with our findings: G3P flux and GPD2 activity were higher in the LT cell lines although there were no differences in glucose flux to lactate between the two species. Compared to the malate-aspartate shuttle the G3P shuttle is less efficient, producing only two molecules of ADP per NADH instead of three, the remaining energy being dissipated as heat (18,21). Suspecting that higher G3P shuttle capacity could be linked to increased thermogenesis in LT cells, we measured changes in intramitochondrial temperature using Mito Thermo Yellow (MTY). MTY is a thermosensitive fluorescent dye which accumulates inside the mitochondrial matrix and changes fluorescence as an inverse function of temperature i.e., higher MTY signal indicates lower mitochondrial temperature (25–27). After inhibition of Complex I by rotenone the drop in mitochondrial temperature (ΔT°) was 35% lower in the LT cells (Figure 2C). In simplified terms, this difference in ΔT° can be explained in two opposite ways: either the temperature of LT mitochondria was lower before the addition of rotenone, or their temperature after rotenone addition dropped less than in LE.

To test which hypothesis is correct and verify the connection between GPD2 activity and mitochondrial energetics, we knocked down GPD2 in the LE and LT fibroblasts. Comparison between *Lepus* and *Homo sapiens* GPD2 consensus sequences allowed the selection of a shRNA from existing human libraries that matched both *Lepus* species as well as the human coding sequence. Silencing GPD2 caused a significant decrease in GPD2 protein level to 60% and 44% of the WT levels respectively in LE3 and LT1 fibroblast cell lines (Figure 2D). In LE3 and LT1 permeabilized fibroblasts, GPD2-KD led to a small decline in respiratory chain activity (Figure S4C), indicating possible changes in OXPHOS function caused by GPD2 silencing. The decrease in GPD2 protein level was associated with a very strong decrease in G3P-driven respiration whether we normalized it to the activity of other OXPHOS complexes or not (Figure 2E, Figures S4D-E). Mitochondrial thermogenic capacity (ϕλT°) of the GPD2-KD fibroblasts was assessed as previously mentioned. It was consistently lower in the GPD2-KD lines (Figure 2F) although the decrease was rather modest (12-27%) compared to GPD2-KD effect on respiration (Figure 2E). This weak effect of GPD2 on mitochondrial thermogenic capacity would suggest it has marginal importance in thermal adaptation and therefore that species difference in ϕλT° are due to the differences in their basal mitochondrial temperature. To assess this inference, we performed comparative measurements of MTY staining using live cells imaging. The LT mitochondria reproducibly and significantly showed higher MTY signal in our cell culture conditions, indicating that they are colder than LE ones (Figure 3A-B). Moreover, GPD2-KD cells showed a small but consistent decrease in their basal mitochondrial temperature somewhat proportional to the decrease in ϕλT° (compare Figure 3B and 2F). Altogether, it suggests that most of the mitochondria from LE and LT species function at different temperatures, which are only marginally influenced by GPD2 activity.

**Figure 3:**
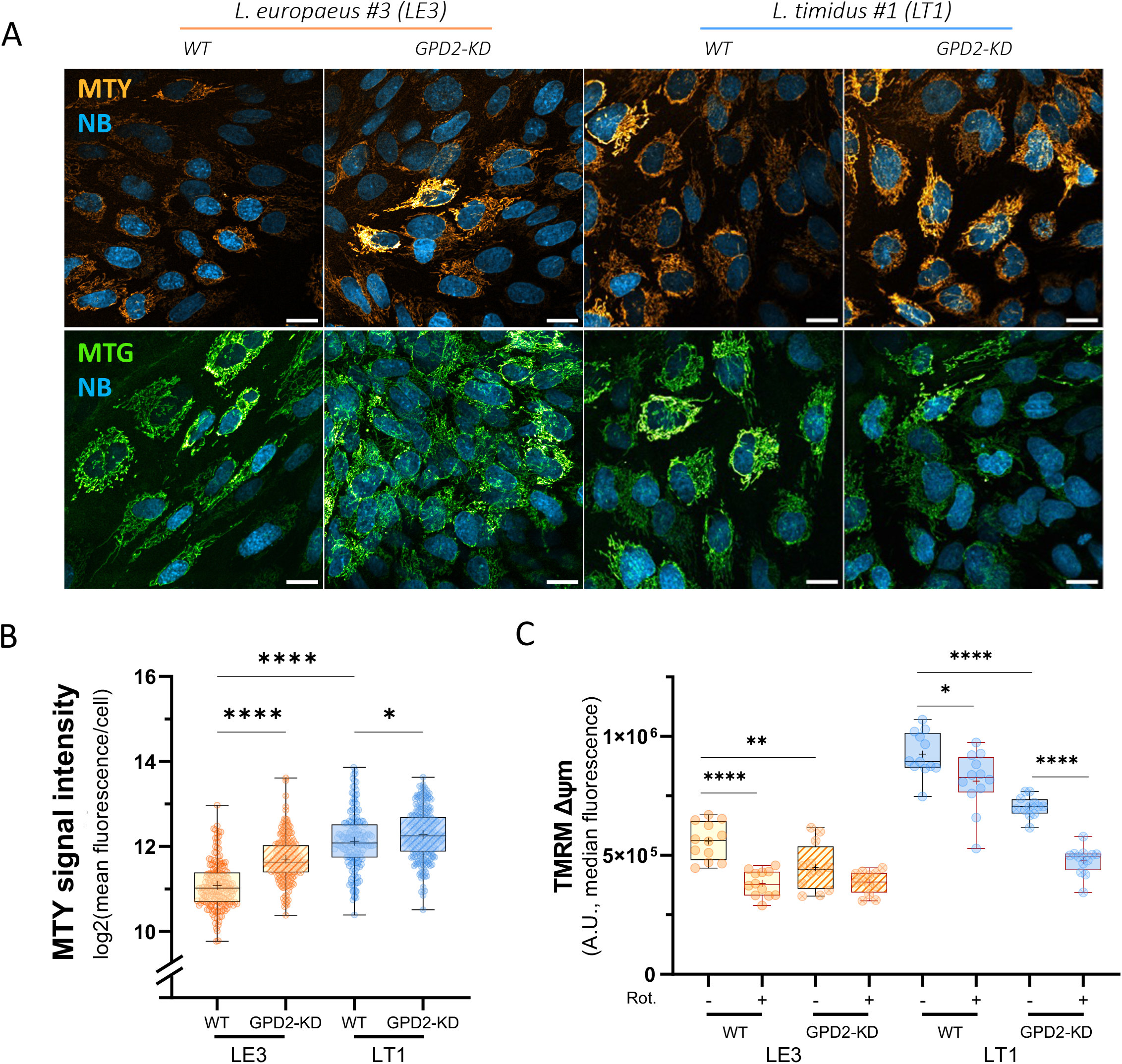
Hare fibroblasts present different basal mitochondrial temperature and membrane potential that are affected by knocking down of GDP2. (A) Visualization of MitoThermo Yellow (MTY) staining in LE and LT WT and GPD2-KD cells by confocal microscopy. Upper row: Intensity of the MTY signal is visually stronger in the GPD2-KD cell lines. Lower row: MitoTracker Green (MTG) staining demonstrates no differences in the mitochondrial network between the cell lines. NB = NucBlue. Scale bar 20 μm. (B) Quantitation of MTY signal from confocal images. LE cells show higher basal mitochondrial temperature compared to LT cells. Knocking down of GDP2 led to lower basal temperature in both species with a more significant drop in LE cells. (C) Quantitation of mitochondrial membrane potential (Δψm) by TMRM. In LE cells both rotenone and GPD2-KD led to similar drop in Δψm. In LT cells, both knock-down of GPD2 and Complex I inhibition were required to drop Δψm at the level of LE cells.

Since GPD2 did not appear critical in establishing basal mitochondrial temperatures, we assessed whether it could have a different species-specific role. Our previous comparative analysis of respiratory chain activity showed a strong difference in mitochondrial membrane potential (ΔΨ_m_) between the two hare species, with the LT fibroblasts having considerably higher ΔΨ_m_ (15). To assess if the mitochondrial membrane potential is connected to the GPD2 activity, we analyzed the effect of GPD2-KD on ΔΨ_m_ by flow cytometry of TMRM stained cells. In accordance with our previous results (15), LT cells maintained ∼40% higher ΔΨ_m_ compared to LE cells and their ΔΨ_m_ was more resilient to Complex I inhibition (Figure 3C) . GPD2 knockdown led to a clear decrease of ΔΨ_m_ in both species (Figure 3C). However significant differences could be observed between species. In LE fibroblasts, the knockdown of GPD2, the abolition of Complex I activity, or a combination of the two led to the same decrease in ΔΨ_m_, suggesting that these activities cannot complement each other, each needing the other to maintain ΔΘ_m_. In contrast, in LT cells abolishing Complex I activity or knocking-down GPD2 caused an intermediate drop in ΔΨ_m_, which was further decreased when the two interventions were combined, suggesting the ΔΨ_m_ effect of one is at least partly independent of the other.

## Discussion

By combining targeted metabolomics, metabolic tracing, high resolution respirometry and subcellular thermometry we have uncovered species-specific metabolic and bioenergetic differences in fibroblasts extracted from two closely related hare species, the mountain hare (*Lepus timidus)* and the brown hare (*Lepus europaeus)*. Mountain hare fibroblasts have higher steady-state levels of G3P and higher flow of glucose towards G3P. This was associated with a stronger G3P-driven respiratory capacity in permeabilized cells. Finally, we demonstrate species-specific difference in how G3P influences mitochondrial membrane potential and to a lesser extent thermal homeostasis. We suggest that these differences reflect, at the cellular level, the environmental pressures and adaptive responses that have shaped the evolution of these two species.

In our previous study (15), we established and characterized a cell culture model, identifying genetic and physiological species differences in fibroblast cell lines extracted from a sympatric population of brown hares and mountain hares in Finland. The most striking outcome of this study was the higher growth rate and wound healing capacity of brown hare cells, which was associated with a lower mitochondrial membrane potential (Δλϑ′_m_). These results suggested species-specific strategies in cellular energy metabolism between the two species, although we could not establish a connection between the mitochondrial and physiological species differences. To address this gap, in the present study, a comparative analysis of the fibroblast metabolome of the two species was established. It resulted in a small number of metabolites (17/143) that were present at significantly different levels. Brown hare cells showed higher levels of amino acid and fatty acid -related metabolites which may be connected to their enhanced proliferation abilities (28), as also observed in our previous study (15). Mountain hare cells present higher levels of G3P, and creatine pathway -related metabolites, both known to be complementary sources of ATP, respectively, from *de novo* mitochondrial production and storage of mitochondrial ATP in the form of PCr. Synthesized from carbohydrates, amino acids, as well as from triglycerides, G3P can be metabolized in the mitochondria by GPD2 to generate ATP and heat, or used for synthesis of glycerolipids, or triglycerides. Increased G3P availability is therefore often seen as a hallmark of improved metabolic flexibility (25). This led us to hypothesize that mountain hare cells might have sacrificed growth in exchange for enhanced metabolic adaptability to external stressors such as cold. Whether the cell lines from the two species differ also in their long-term energy storage e.g., lipids, remains to be studied.

Metabolic tracing of ^13^C glucose showed a great proportion of the carbons being directed to lactate in both species. This glycolytic switch is a classic and reversible adaptive change in cells maintained under high glucose conditions, especially in fibroblasts (29). Despite this metabolic reprogramming in cells from both species, the flux of glucose to G3P was significantly higher in LT fibroblasts. In addition, the proportion of labelled dihydroxyacetone-phosphate (DHAP), the direct precursor of G3P, was also higher in LT fibroblasts, although at a later timepoint. This is consistent with the activation of the G3P shuttle, where cytosolic NADH is used by the GPD1 to convert DHAP originating from ^13^C glucose metabolism into G3P. G3P is then metabolized by the mitochondrial GPD2 returning to the cytosol as DHAP. Assuming that GPD2 activity is the rate limiting G3P shuttle reaction, activation of this cycle should lead to the successive accumulation of G3P and then DHAP, as we see in LT fibroblasts. Indeed, the G3P shuttle has been described as limited by GPD2 protein levels and allosteric regulations in mammalian tissues (18). GPD2 regulation is complex, being activated by cytosolic Ca^2+^ and to a lesser extent other divalent cations, while inhibited by glyceraldehyde 3-phosphate, palmitoyl-CoA and free fatty acids. Since western blot analyses showed no difference in GPD2 protein level between the two species, one would privilege the second hypothesis. However, GPD2 sequence differs between the LT and LE species at two amino acid positions, one of which comparatively decreases *in silico* matching of the LE sequence with the consensus sequence for FAD dependent oxidoreductases. It is possible that the differences in protein activity originate from the species differences in their sequences. Recently, the human GPD2 was purified and characterized *in vitro* (30) and a similar procedure could be used to compare the catalytic properties but also allosteric regulation of the brown and mountain hare enzymes.

G3P-driven respiration was significantly higher in mountain hare cells which confirmed that the two species have different GPD2 capacity. GPD2 is the rate limiting step in the G3P shuttle which participates in the reoxidation of cytosolic NADH and the regulation of NADH/NAD ratio (18). Two major mechanisms, the malate-aspartate- and the G3P shuttles can transfer NADH reducing power from the cytosol to the mitochondria, where it fuels ATP production by the OXPHOS system. Unlike the malate-aspartate shuttle which allows a net transfer of cytosolic NADH to the Complex I, the G3P shuttle branches on the quinone downstream of Complex I, releasing the energy which would have been used to pump protons at Complex I as heat (18).

MTY thermometry reliably assesses the relative (before *vs* after treatment) temperature of mitochondria (31,32). As mitochondrial temperature can drop to that of their environment, (37°C in mammalian cells), but not further, one can also infer the absolute temperature of mitochondria before treatment. For example, a drop of 15°C in temperature induced by rotenone treatment would imply that mitochondrial temperature was at least 52°C before the treatment. This method led to the finding that mitochondrial temperature of human fibroblasts was ≥ 50°C (31), an estimation which was confirmed in various human and animal cell types (32). With the same approach we observed a drop of ∼15°C and ∼10°C, respectively, in Complex I inhibited mitochondria from brown and mountain hare fibroblasts, suggesting initial temperatures of ≥ 52°C and ≥ 47°C. These estimates are in good agreement with previous knowledge from other cell models (31,32). Alternatively, the activity of the G3P shuttle, which is theoretically independent of Complex I, could preserve mitochondrial temperature in mountain hare, limiting the drop induced by Complex I inhibition.

Knocking down GPD2 strongly decreased G3P shuttle activity in permeabilized cells but only slightly lowered basal mitochondrial temperature. If GPD2 activity is thermogenic, as previously shown (33), why does it have so little impact on mitochondrial temperature despite significantly lower activity? In the reported studies, GPD2 effects on temperature are measured in whole animals or cells. The MTY thermometric dye is located in the mitochondrial matrix, thereby detecting intramitochondrial temperature. As GPD2 is sitting inside the intermembrane space, the heat produced by the activity of the enzyme may radiate outside of mitochondria. Such thermogenic directionality could be dependant on properties of GPD2 itself or its environment, including the structure and composition of the inner and outer mitochondrial membranes. Further explorations of this theory would require the use of thermosensitive intermembrane space reporters, or to compare the temperature in the vicinity of the inner and outer mitochondrial membranes.

In contrast to its species-independent effect on mitochondrial temperature, GPD2-KD had strikingly different impact on mitochondrial membrane potential. In LE fibroblasts, either Complex I inhibition or knocking down of GPD2 was enough to suppress ΔΨ_m_ efficiently, while in LT fibroblasts, the combination of the two was needed to obtain the same effect. The associations of GPD2 with other mitochondrial membrane proteins and higher assemblies including OXPHOS complexes has been speculated but is yet to be shown (18). Formation of supercomplexes is presumed to provide for example more efficient substrate channeling and minimize electron leak. Reverse activity of the ATP synthase where the complex consumes ATP to pump H^+^ outside of the mitochondrial matrix, is known to participate in ΔΨ_m_ maintenance, particularly in situations of OXPHOS deficiency (34). Species-specific difference in the reverse flow through ATP synthase may explain in part the effect of GPD-KD on ΔΨ_m_. Different effects of GPD2-KD on the ΔΨm in the two hare species point to different associations with at least Complex I. Complex I and high ΔΨm are both strongly linked to ROS-production (18). Variation in GPD2-related ROS production has been found between tissues, the significance of which seems to be independent of GPD2 expression, further supporting the idea of different associations with the OXPHOS complexes (18,35).

## Conclusions

Our study shows divergent metabolic strategies between fibroblasts extracted from two evolutionary distinct hare species, the European brown hare (*Lepus europaeus,* LE) and the mountain hare (*Lepus timidus,* LT). One of the main findings is the difference in glycerol 3-phosphate (G3P) metabolism and activity of the mitochondrial glycerol 3-phosphate dehydrogenase (GPD2). This is also the first study to present variation in thermogenic properties of skin-derived fibroblasts, with LE fibroblasts showing higher basal mitochondrial temperature compared to LT. The mechanisms behind the different role of GPD2 activity in the two species remains to be studied but it is suggested to be due to differences in the structure of the active site and/or associations between the enzyme and the OXPHOS complexes such as Complex I. Unveiling these mechanisms may help to understand the evolution of cellular metabolic traits and more importantly, their plasticity and predict how they may advance or hamper the survival of the organism in a stochastically changing climate.

## Materials and Methods

### Ethics statement

The study did not involve experimentation on live animals. All experiments were performed using cultured cell lines. Both species of hare are legal game animals in Finland and the sampling was previously described (15) hence no ethical assessment was required.

### Cell lines and cell culture

Four immortalized mountain hare (LT 1, 4, 5, 6) and brown hare (LE 1, 2, 3, 4) fibroblast cell lines were generated through SV40 transformation as previously described (15). The chromosomal integrity and the gene expression of the immortalized fibroblast was verified using high throughput sequencing and transcriptomics, none of the cell lines presented genetic abnormality and their transcriptome signature remained that of skin fibroblasts (36). Cells were grown in standard cell culture conditions (37 °C, 5 % CO2, ∼95 % relative humidity). DMEM-Hi glucose growth medium (# D6546) supplemented with 10 % (vol/vol) heat-inactivated fetal bovine serum (HI-FBS, # F7524), 1 % L-glutamine (# G7513, Sigma-Aldrich), and 1 % penicillin/streptomycin (P/S, # 15140122, Gibco) was used to culture the cells. Stable glycerol-3-phosphate dehydrogenase 2 (GPD2) knock-down (KD) cell lines were generated from parental mountain hare (LT1) and brown hare (LE3) cell lines by lentiviral transduction of shRNA (TRCN0000028617 and TRCN0000028579 MISSION^®^ shRNAs, Sigma-Aldrich) followed by puromycin-resistant marker selection.

### Immunoblotting

Cells were tested for the expression of GPD2 protein using Western blot as previously described (15). Briefly, protein samples (30 μg) were loaded on 4 - 20 % gels (# 4561093, Bio-Rad), run at 100 V and transferred to nitrocellulose membranes (# 1704158, Bio-Rad). Blots were blocked in 5 % milk diluted in tris buffered saline plus 0.1 % Tween 20 (TBS-T) and probed with GPD2 polyclonal antibody (# 17219-1-AP, Proteintech) diluted 1:2000 in blocking solution at 4 °C overnight followed by incubation with peroxidase-linked goat anti-rabbit IgG (1:10000, # PI-1000-1, Vector Laboratories). Chemiluminescent substrate (# 34095, Thermo Scientific) was applied to visualize immunoreactive bands in ChemiDoc XRS+ Imaging System (Bio-Rad). Blots were washed with TBS-T, re-probed with mouse monoclonal anti-Vinculin antibody (1:10000, # V9264, Sigma-Aldrich) followed by peroxidase-conjugated horse anti-mouse IgG (1:10000, # PI-2000, Vector Laboratories) and imaged as described above. Images were captured and analysed using Image Lab 6.0.1 software. GPD2 protein expressions were normalized to Vinculin. Three individual experiments were performed for each cell line, with additional two experiments for LT1, LE3 and for the established GPD2-KD cell lines.

### Flow cytometry

Cells were collected by trypsinization, pelleted at 250 g for 3 min and resuspended with DPBS (# D8537, Sigma-Aldrich) at 1x10^6^ cells/ml. All samples, excluding a negative control, were stained with 20 nM TMRM (# T668, Invitrogen), a mitochondrial dye used to estimate mitochondrial membrane potential (Δψm), in the presence of 50 µM Verapamil (# V4629, Sigma-Aldrich), an efflux pump inhibitor (37) for 20 min at 37 °C. Half of the samples were treated with 0.075 µM rotenone (# R8875; Sigma-Aldrich), a mitochondrial Complex I inhibitor. After incubation, cells were pelleted and washed with DPBS twice, resuspended with DPBS containing 2 % FBS, and median fluorescent intensities of TMRM (Ex 561 nm, Em 584) were measured with CytoFlex S flow cytometer (Beckman Coulter). For each studied cell line four independent experiments with three technical repeats were performed as described above. Data were analysed using BD FlowJo 10.7.2 software.

### High-resolution respirometry

Cells were grown until 80 % confluence and collected by trypsinization. For each measurement, 5×10^6^ cells were resuspended in the respiration buffer (225 mM sucrose; 75 mM mannitol; 10 mM KCl; 10 mM KH2PO4; 5 mM MgCl2; 10 mM Tris base; 1 mg/ml BSA; pH 7.4) and transferred into the oxygen-calibrated chamber of Oxytherm respirometer (Hansatech Instruments). Cell membranes were permeabilised with 55 µM digitonin. Mitochondrial respiration was assessed by additions of substrates and inhibitors in the following order: pyruvate + glutamate + malate (5 mM each; # P8574, # G5889, # M7397; Sigma-Aldrich), ADP (1 mM; # 117105; Calbiochem), rotenone (0.15 µM; # R8875; Sigma-Aldrich), glycerol 3- phosphate, G3P (10 mM; # 94124; Sigma-Aldrich), antimycin A (90 ng/ml; # A8674; Sigma-Aldrich), ascorbate (700 µM; # A4034; Sigma-Aldrich), N,N,N′,N′-tetramethyl- p-phenylenediamine (300 µM, # T7394; Sigma-Aldrich) and potassium cyanide (200 μM; # 60178; Sigma-Aldrich). Oxygen consumption (pmol.sec−1.ml−1) was normalized to the cell count (i.e., per million of cells) as well as to the Complex I and Complex IV driven respiration. For each cell line, seven independent experiments were performed as described above. Statistics were performed using a nested glm (response variable ∼ species/cell line) and fitting a gamma distribution. When only one cell line was included per species, a one-way anova was preferred.

### Spectrofluorometry

Mitochondrial temperature was measured using spectrofluorometry as previously described (36). Briefly, cells were stained for 15 min at 37 °C with 100 nM Mito Thermo Yellow (MTY), a thermosensitive fluorescent probe (38), trypsinized and washed with DPBS. Around 5x10^6^ cells resuspended in DPBS were transferred into a quartz cuvette with 10 mm optical path (Hellma Analytics, Germany). The cuvette was placed in a Peltier chamber pre-set to 38 °C that allows for precise and rapid temperature control inside a fluorometer system (PTI QuantaMasterTM, LPS-220B, Horiba). Kinetic measurements of MTY fluorescence signal (Ex 542 nm, Em 562 nm) were collected using FelixGX software. Once the fluorescence signal stabilized, the cells were treated with 0.075 µM rotenone. After the fluorescence signal reached a steady state, the temperature was changed to 41 °C, and then back to 38 °C, which allowed for MTY fluorescence signal calibration. Three independent experiments were performed for each cell line, with additional two experiments for LT1, LE3 and for the established GPD2-KD cell lines.

### Imaging

Cells were seeded on poly-d-lysine coated glass coverslips (# P35GC-1.5-14-C, MatTek) and grown in standard cell culture conditions for 48 h. Cell nuclei were stained with NucBlue (NB, # R37605, Invitrogen) as specified by manufacturer, and mitochondria with either 100 nM MTY for 15 min or 100 nM MitoTracker Green FM (MTG, # M7514, Invitrogen) for 30 min at 37 °C. Stained cells were washed with DPBS and maintained in FluoroBrite DMEM (# A1896701, Gibco) supplemented with 1% P/S to preserve cell fitness throughout the imaging process. Culture dishes were placed inside the microscope onstage incubator pre-set to maintain standard cell culture conditions during live-cell imaging. Relative z-stacks of cells stained with NB (Ex 405, Em 450/50), MTY (Ex 488, Em 540/30) and MTG (Ex 488, Em 525/50) were acquired from five non-overlapping imaging fields using Nikon’s A1R+ confocal laser scanning microscope system (Nikon Eclipse Ti2-E), Perfect Focus System® and Nikon 60x / 1.27 water-immersion objective (CFI SR Plan Apo IR 60XC WI). The detector (A1- DUG GaAsP Multi Detector Unit) settings were adjusted for each channel to avoid image saturation and kept constant for all experiments. Images were deconvolved using Huygens Essential (SVI; Hilversum) and analyzed in Fiji-ImageJ (39). Midsections from acquired z-stacks were used to measure mean MTY signal intensities of individual cells. For visualization purpose, SD z-projections were created, background was subtracted, brightness and contrast were adjusted. Adjustment settings were kept constant to obtain representative figures.

### Metabolic phenotyping

All cell lines (4 LT and 4 LE lines) were collected by trypsinization, counted with EVE™ automatic cell counter (NanoEntek) and inoculated in IF-M2 medium (# 72302, Biolog) supplemented with 5% HI-FBS, 1% P/S and 0.3 mM L-glutamine. Cells were seeded at density 2x10^4^ cells/well on 96-well plates containing carbohydrate/carboxylate substrates (PM-M1, # 13101, Biolog) and cultured under standard cell culture conditions for 48 h. A redox dye mix MB (# 74352, Biolog) was added into each well at 10 µl/well. Absorbance of NADH signal accumulation (562 nm) was measured with pre-warmed to 37°C Elmer EnVision 2104 plate reader (Perkin) immediately after adding the dye (0 h) and at time points 3 h, 7 h, 21 h, 27 h, 48 h, 54 h. Between the measurements, cells were maintained in the incubator under standard cell culture conditions. Absorbance values above 4 were regarded as aberrant and removed. Comparison of substrate usage between hare species was done in R (version 4.3.1) using opm package (40). Modelling of NADH accumulation was obtained by fitting an exponential plateau model (〖Nadh〗_((t))=〖Nadh〗_max-(〖Nadh〗_max-〖Nadh 〗_0) e^(-kt)). We compared maximum NADH levels (〖Nadh〗_max), controlling for family-wise error-rate to measure the differences in NADH production. For each timepoint and species, the signal from each well was averaged to obtain (Lt_m) ̅ and (Le_m) ̅. For each metabolite (m) and timepoint, relative differences ((Le_m) ̅- (Lt_m) ̅) / 〖((Le;Lt) ̅)〗_m were calculated, averaged and plotted.

### Metabolomics

All cell lines were seeded 6-well plates and grown for 3 days till reaching 80 % confluence. The medium was changed 4 h before polar metabolites extraction. The plates were removed from the incubator and put on dry ice. Cells were rinsed with ice-cold 150 mM NH4AcO (pH 7.3) and incubated with 1 ml/well 80 % MeOH at -80 °C for 45 min to aid quenching and protein precipitation. Cells were scraped from individual wells and collected in Eppendorf tubes (n = 6). Samples were vortexed and centrifuged at 1.6x10^4^ g for 15 min at 4 °C. Supernatants were transferred into fresh Eppendorf tubes, dried using vacuum concentrator (ScanVac MaxiVac, LaboGene) and stored at -80 °C. Cell pellets were lysed in 0.1M NaOH, and protein concentrations were estimated using BCA assay kit (# 23225, Thermo Scientific) as specified by manufacturer. Samples were sent for targeted analysis to UCLA Metabolomics Center (Los Angeles, USA). Samples 42 (LT5), 44 and 46 (LT6) were excluded due chromatographic aberrations (i.e., metabolite retention shift). Eleven metabolites with less than 50% of non-zero measures per group were excluded from the analysis. For each sample metabolite levels were normalized by protein content. Metabolites were either compared using Student’s T-test with no FDR correction and no normalization (Figure 1A); or normalized by the median, auto-scaled, and compared using SAM (Significance Analysis of Metabolomics) with FDR correction (p < 0,05)(41). SAM analysis identified 6 significantly elevated metabolites in LT (same as in Figure 1A excluding homocysteine) and 14 metabolites with significantly lower levels (same as in Figure 1A excluding deoxycytosine and including HMG-CoA, carbamoyl-aspartate, 4-OH-phenyllactate, sarcosine, glutamate and aspartate).

### 13C-glucose tracing

Two mountain hare cell lines (LT 5 and 6) and two brown hare (LE 3 and 4) were seeded on 6-well plates and grown for 3 days till reaching 80 % confluency. Before extraction, the medium was changed to DMEM without glucose (# 01-057-1A Sartorius) supplemented with 25 mM final concentration of ^13^C-glucose (# CLM-1396- 1, Cambridge Isotope Laboratories). After 30 min and 120 min exposures, the metabolites were extracted as described above. Samples were sent for flux analysis to FIMM Metabolomics Unit at University of Helsinki (Helsinki, Finland). For each isotopologue, the levels were divided by the total amount of labelled metabolites detected in the sample at the given time. The variation in this proportion were then analyzed in R using a linear mixed model: lmer(metabolite ∼ Timept * Genotype + (1|strain)). Note that fluxes of ^13^C-glucose to phosphoserine were not analyzed due to a very low number of reliable measures.

### Transcriptomics

We pooled all genes having GO-terms related to 3P with those as involved in Glo3P metabolism from the KEGG library (n=146) and did the same for the genes related to Cr and its metabolic pathway (n=13). The transcriptomes of the LT and LE cells have been published earlier (19). The raw sequence reads are available through BioProject accession PRJNA826339.

### Protein sequence

Sequence alignments were perform using ApE v3.1.4. Comparison with known consensus sequences were performed using MyHits (http://myhits.isb-sib.ch) (42).

### Statistics

Before analyses, normality and lognormality tests were used (i.e., d’Agostini & Pearson and Shapiro-Wilk). Normal and lognormal distributions were compared using a maximum likelihood method and generated QQ plots. If lognormal distribution was more likely, the data was log2 transformed and statistical analysis was done on logs. Possible outliers were identified and removed using Grubb’s (alpha = 0.2) or ROUT (FDR 2 %) methods depending on the number of expected outliers. For comparisons between two groups (i.e., species), unpaired Student’s t test (assuming equal variance) was performed. For comparisons between multiple groups, one- or two- way ANOVA was performed with subsequent Tukey’s multiple comparisons test, and adjusted p values were reported. Statistical tests were conducted at a 5% significance level. GraphPad Prism 9.0.0 was used for statistical analyses and graphical representation of the obtained results if not otherwise stated.

Analyses of the respirometry and ^13^C-glucose tracing data were performed using R 4.3 in RStudio 2023.03.1.0. Species was set as a predictor of interest, in which cell line was set a nested factor. The effect response variable was analyzed using glm or lmer as presented in the corresponding methods. To identify the best type of distribution, we used gamlss 5.4.12 library of distributions from the Generalized Additive Models for Location Scale and Shape (43). The distributions were ranked using corrected generalized Akaike Information Criteria (AIC). For each data set, the distribution with the lowest AIC score compatible with glm or lmer was selected. The the importance of each predictor and eventual interaction was assessed using ANOVA. Key results are reported in the figure legends (*P < 0.05; **P < 0.01; ***P < 0.001; ****P < 0.0001; ns, non-significant).

## Acknowledgements

Authors acknowledge the support provided by Johanna ten Hoeve at UCLA Metabolomics center, US; Anni I. Nieminen at Biocenter Finland and HiLIFE-funded FIMM Metabolomics Unit, Helsinki, Finland; Young-Tae Chang at POSTECH, Republic of Korea, for providing MTY dye; Howy Jacobs for invaluable feedback and insightful discussions; Teemu O. Ihalainen at Tampere Imaging Facility (TIF), Tampere University, Finland for helping with the imaging.

## Abbreviations

Δψm: mitochondrial membrane potential
ATP: adenosine triphosphate
BSA: bovine serum albumin
CDS: coding DNA sequence
Cr: creatine
DMEM: Dulbecco’s Modified Eagle Medium
DPBS: Dulbecco’s phosphate-buffered saline
ECM: extracellular matrix
ex: excitation
em: emission
FA: fatty acids
FCCP: carbonyl cyanide-p-trifluoromethoxyphenylhydrazone
G3P: glycerol-3-phosphate
GPD2: mitochondrial glycerol-3-phosphate dehydrogenase
h: hours
HI-FBS: heat inactivated fetal bovine serum
min: minutes
MTG: Mito Tracker Green
MTT: methylthiazolyldiphenyl-tetrazolium bromide
MTY: Mito thermo yellow
NB: NucBlue
PCr: phosphocreatine
RT: room temperature
ROI: region of interest
SRB: sulforhodamine B sodium salt
TMRM: tetramethylrhodamine methyl ester
TBS-T: tris buffered saline plus 0.1% Tween 20
TCA: tricarboxylic acids cycle
P/S: penicillin/streptomycin

## Data Availability

All relevant data are within the paper and its Supplementary Information files.

## Funding

S.S. was funded by AFM Telethon (23527), Alfred Kordelin Foundation (200340) and Tampere Institute of Advanced Studies. E.D. and J.P. were funded by the Academy of Finland (R’Life 329264). The funders had no role in study design, data collection and analysis, decision to publish, or preparation of the manuscript.

**List of supplementary files** Supplementary table S1 Supplementary table S2 Supplementary table S3

## References

1. Talbott HE, Mascharak S, Griffin M, Wan DC, Longaker MT. Wound healing, fibroblast heterogeneity, and fibrosis. Cell Stem Cell. 2022 Aug;29(8):1161–80.

2. Unger C, Felldin U, Rodin S, Nordenskjöld A, Dilber S, Hovatta O. Derivation of Human Skin Fibroblast Lines for Feeder Cells of Human Embryonic Stem Cells. Curr Protoc Stem Cell Biol [Internet]. 2016 Feb [cited 2024 Jan 15];36(1). Available from: https://currentprotocols.onlinelibrary.wiley.com/doi/10.1002/9780470151808.sc01c07s36

3. Lendahl U, Muhl L, Betsholtz C. Identification, discrimination and heterogeneity of fibroblasts. Nat Commun. 2022 Jun 14;13(1):3409.

4. Shani O, Raz Y, Monteran L, Scharff Y, Levi-Galibov O, Megides O, et al. Evolution of fibroblasts in the lung metastatic microenvironment is driven by stage-specific transcriptional plasticity. eLife. 2021 Jun 25;10:e60745.

5. Harper JM, Wang M, Galecki AT, Ro J, Williams JB, Miller RA. Fibroblasts from long-lived bird species are resistant to multiple forms of stress. J Exp Biol. 2011 Jun 1;214(11):1902–10.

6. Jimenez AG, Harper JM, Queenborough SA, Williams JB. Linkages between the life-history evolution of tropical and temperate birds and the resistance of their cells to oxidative and non- oxidative chemical injury. J Exp Biol. 2012 Jan 1;jeb.079889.

7. Harper JM, Salmon AB, Leiser SF, Galecki AT, Miller RA. Skin-derived fibroblasts from long-lived species are resistant to some, but not all, lethal stresses and to the mitochondrial inhibitor rotenone. Aging Cell. 2007 Feb;6(1):1–13.

8. Salmon AB, Akha AAS, Buffenstein R, Miller RA. Fibroblasts From Naked Mole-Rats Are Resistant to Multiple Forms of Cell Injury, But Sensitive to Peroxide, Ultraviolet Light, and Endoplasmic Reticulum Stress. J Gerontol A Biol Sci Med Sci. 2008 Mar 1;63(3):232–41.

9. Norin T, Metcalfe NB. Ecological and evolutionary consequences of metabolic rate plasticity in response to environmental change. Philos Trans R Soc B Biol Sci. 2019 Mar 18;374(1768):20180180.

10. Levänen R, Thulin CG, Spong G, Pohjoismäki JLO. Widespread introgression of mountain hare genes into Fennoscandian brown hare populations. Castiglia R, editor. PLOS ONE. 2018 Jan 25;13(1):e0191790.

11. Thulin C. The distribution of mountain hares *Lepus timidus* in Europe: a challenge from brown hares *L. europaeus* ? Mammal Rev. 2003 Mar;33(1):29–42.

12. Angerbjorn A, Flux JEC. Lepus timidus. Mamm Species. 1995 Jun 23;(495):1.

13. Bock A. *Lepus europaeus* (Lagomorpha: Leporidae). Harris JM, Hamilton MJ, editors. Mamm Species. 2020 Dec 23;52(997):125–42.

14. Pohjoismäki JLO, Michell C, Levänen R, Smith S. Hybridization with mountain hares increases the functional allelic repertoire in brown hares. Sci Rep. 2021 Aug 4;11(1):15771.

15. Gaertner K, Michell C, Tapanainen R, Goffart S, Saari S, Soininmäki M, et al. Molecular phenotyping uncovers differences in basic housekeeping functions among closely related species of hares (*Lepus* spp., Lagomorpha: Leporidae). Mol Ecol. 2023 Aug;32(15):4097–117.

16. Martinez-Reyes I, Chandel NS. Mitochondrial TCA cycle metabolites control physiology and disease. Nat Commun. 2020 Jan 3;11(1):102.

17. Kazak L, Rahbani JF, Samborska B, Lu GZ, Jedrychowski MP, Lajoie M, et al. Ablation of adipocyte creatine transport impairs thermogenesis and causes diet-induced obesity. Nat Metab. 2019 Feb 25;1(3):360–70.

18. Mráček T, Drahota Z, Houštěk J. The function and the role of the mitochondrial glycerol-3- phosphate dehydrogenase in mammalian tissues. Biochim Biophys Acta BBA - Bioenerg. 2013 Mar;1827(3):401–10.

19. Wyss M, Kaddurah-Daouk R. Creatine and Creatinine Metabolism. Physiol Rev. 2000 Jul 1;80(3):1107–213.

20. Kazak L, Cohen P. Creatine metabolism: energy homeostasis, immunity and cancer biology. Nat Rev Endocrinol. 2020 Aug;16(8):421–36.

21. Beignon F, Gueguen N, Tricoire-Leignel H, Mattei C, Lenaers G. The multiple facets of mitochondrial regulations controlling cellular thermogenesis. Cell Mol Life Sci. 2022 Oct;79(10):525.

22. Bochner BR, Siri M, Huang RH, Noble S, Lei XH, Clemons PA, et al. Assay of the Multiple Energy- Producing Pathways of Mammalian Cells. Tomé D, editor. PLoS ONE. 2011 Mar 24;6(3):e18147.

23. Žigon P, Mrak-Poljšak K, Lakota K, Terčelj M, Čučnik S, Tomsic M, et al. Metabolic fingerprints of human primary endothelial and fibroblast cells. Metabolomics. 2016 May;12(5):92.

24. Chretien D, Rustin P, Bourgeron T, Rötig A, Saudubray JM, Munnich A. Reference charts for respiratory chain activities in human tissues. Clin Chim Acta. 1994 Jul;228(1):53–70.

25. Dhoundiyal A, Goeschl V, Boehm S, Kubista H, Hotka M. Glycerol-3-Phosphate Shuttle Is a Backup System Securing Metabolic Flexibility in Neurons. J Neurosci. 2022 Sep 28;42(39):7339–54.

26. Sun Y, Rahbani JF, Jedrychowski MP, Riley CL, Vidoni S, Bogoslavski D, et al. Mitochondrial TNAP controls thermogenesis by hydrolysis of phosphocreatine. Nature. 2021 May 27;593(7860):580– 5.

27. Russell AP, Ghobrial L, Wright CR, Lamon S, Brown EL, Kon M, et al. Creatine transporter (SLC6A8) knockout mice display an increased capacity for in vitro creatine biosynthesis in skeletal muscle. Front Physiol [Internet]. 2014 Aug 26 [cited 2024 Jan 15];5. Available from: http://journal.frontiersin.org/article/10.3389/fphys.2014.00314/abstract

28. Currie E, Schulze A, Zechner R, Walther TC, Farese RV. Cellular Fatty Acid Metabolism and Cancer. Cell Metab. 2013 Aug;18(2):153–61.

29. Lemons JMS, Feng XJ, Bennett BD, Legesse-Miller A, Johnson EL, Raitman I, et al. Quiescent Fibroblasts Exhibit High Metabolic Activity. Goodell MA, editor. PLoS Biol. 2010 Oct 19;8(10):e1000514.

30. Syngkli S, Das B. Purification and characterization of human glycerol 3-phosphate dehydrogenases (mitochondrial and cytosolic) by NAD+/NADH redox method. Biochimie. 2023 Nov;214:199–215.

31. Chrétien D, Bénit P, Ha HH, Keipert S, El-Khoury R, Chang YT, et al. Mitochondria are physiologically maintained at close to 50 °C. Lane N, editor. PLOS Biol. 2018 Jan 25;16(1):e2003992.

32. Terzioglu M, Veeroja K, Montonen T, Ihalainen TO, Salminen TS, Bénit P, et al. Mitochondrial temperature homeostasis resists external metabolic stresses. eLife. 2023 Dec 13;12:RP89232.

33. Silva JE. Thermogenic Mechanisms and Their Hormonal Regulation. Physiol Rev. 2006 Apr;86(2):435–64.

34. Acin-Perez R, Benincá C, Fernandez Del Rio L, Shu C, Baghdasarian S, Zanette V, et al. Inhibition of ATP synthase reverse activity restores energy homeostasis in mitochondrial pathologies. EMBO J. 2023 May 15;42(10):e111699.

35. Mráček T, Pecinová A, Vrbacký M, Drahota Z, Houštěk J. High efficiency of ROS production by glycerophosphate dehydrogenase in mammalian mitochondria. Arch Biochem Biophys. 2009 Jan;481(1):30–6.

36. Gaertner K, Michell C, Tapanainen R, Goffart S, Saari S, Soininmäki M, et al. Molecular phenotyping uncovers differences in basic housekeeping functions among closely related species of hares (*Lepus* spp., Lagomorpha: Leporidae). Mol Ecol. 2022 Nov;mec.16755.

37. Morganti C, Bonora M, Ito K, Ito K. Electron transport chain complex II sustains high mitochondrial membrane potential in hematopoietic stem and progenitor cells. Stem Cell Res. 2019 Oct;40:101573.

38. Arai S, Suzuki M, Park SJ, Yoo JS, Wang L, Kang NY, et al. Mitochondria-targeted fluorescent thermometer monitors intracellular temperature gradient. Chem Commun. 2015;51(38):8044–7.

39. Schindelin J, Arganda-Carreras I, Frise E, Kaynig V, Longair M, Pietzsch T, et al. Fiji: an open-source platform for biological-image analysis. Nat Methods. 2012 Jul;9(7):676–82.

40. Vaas LAI, Sikorski J, Hofner B, Fiebig A, Buddruhs N, Klenk HP, et al. opm: an R package for analysing OmniLog® phenotype microarray data. Bioinformatics. 2013 Jul 15;29(14):1823–4.

41. Xia J, Psychogios N, Young N, Wishart DS. MetaboAnalyst: a web server for metabolomic data analysis and interpretation. Nucleic Acids Res. 2009 Jul 1;37(Web Server):W652–60.

42. Pagni M, Ioannidis V, Cerutti L, Zahn-Zabal M, Jongeneel CV, Hau J, et al. MyHits: improvements to an interactive resource for analyzing protein sequences. Nucleic Acids Res. 2007 May 8;35(Web Server):W433–7.

43. Stasinopoulos DM, Rigby RA. Generalized Additive Models for Location Scale and Shape (GAMLSS) in R. J Stat Softw [Internet]. 2007 [cited 2024 Jan 15];23(7). Available from: http://www.jstatsoft.org/v23/i07/

