## Supplementary figures for "A species difference in glycerol-3-phosphate metabolism reveals metabolic adaptations in hares"

**Supplementary Table and Figure Legends**

***Supplementary Table S1. Targeted metabolomics.*** *Identified metabolites are presented in rows, with their corresponding KEGG ID. Samples are labelled ED.001 to ED.048, followed by the identity of the species (LE or LT) and the cell line number. Metabolite intensity values are presented after normalization to protein content. The values of "between species" F Test and Student T Test are presented in the two last columns.*

***Supplementary Table S2. Biolog Phenotype MicroArrayTM analysis.*** *The first sheet presents the layout of the 96 well plate, indicating which metabolite is present in which well. The second sheet (data) present the absorbance measurement for each metabolite (columns) in function of the cell lines * timepoint (rows).*

***Supplementary Figure S1. Transcriptomics analysis of the creatine pathway - related genes.*** *Simplified map of the metabolic pathway centered around Creatine. Mitochondrial glycine amidinotransferase (GATM) responsible for synthesizing the precursor for creatine, and two alkaline phosphatases converting PCr to creatine + P were only found in the LT cells (in blue).*

***Supplementary Figure S2. Supplementary data on fluxomics analysis of $^{13}C$ glucose.*** *Graphs shows proportion of $^{13}C$ glucose converted into indicated metabolites in LT (red) and LE (blue) cell lines. DHAP = Dihydroxyacetone phosphate, Ribose 5P = Ribose 5-phosphate, Glycerol 3P= Glycerol 3-phosphate.*

***Supplementary Figure S3. GPD2 protein sequence comparison between LE and LT.*** *(A) Protein sequences of GPD2 between the two hare species differ by two amino acids. (B) In LE, a serine (S) is present at position 187, In LT an asparagine (N) is present at the same position.  This serine 187 increase the overall divergence from the consensus motif for FAD dependent oxidoreductases (compare the E-values).*

***Supplementary Figure S4. Supplementary data on high-resolution respirometry.*** *(A) GPD2-driven respiration normalized to complex IV activity. (B) GPD2-driven respiration without normalization. (C) Effect of GPD2-KD on complex I -driven respiration. (D) Effect of GPD2-KD on GPD2-driven respiration normalized to complex IV activity. (E) Effect of GPD2-KD on GPD2-driven respiration without normalization.*

### Figure S1

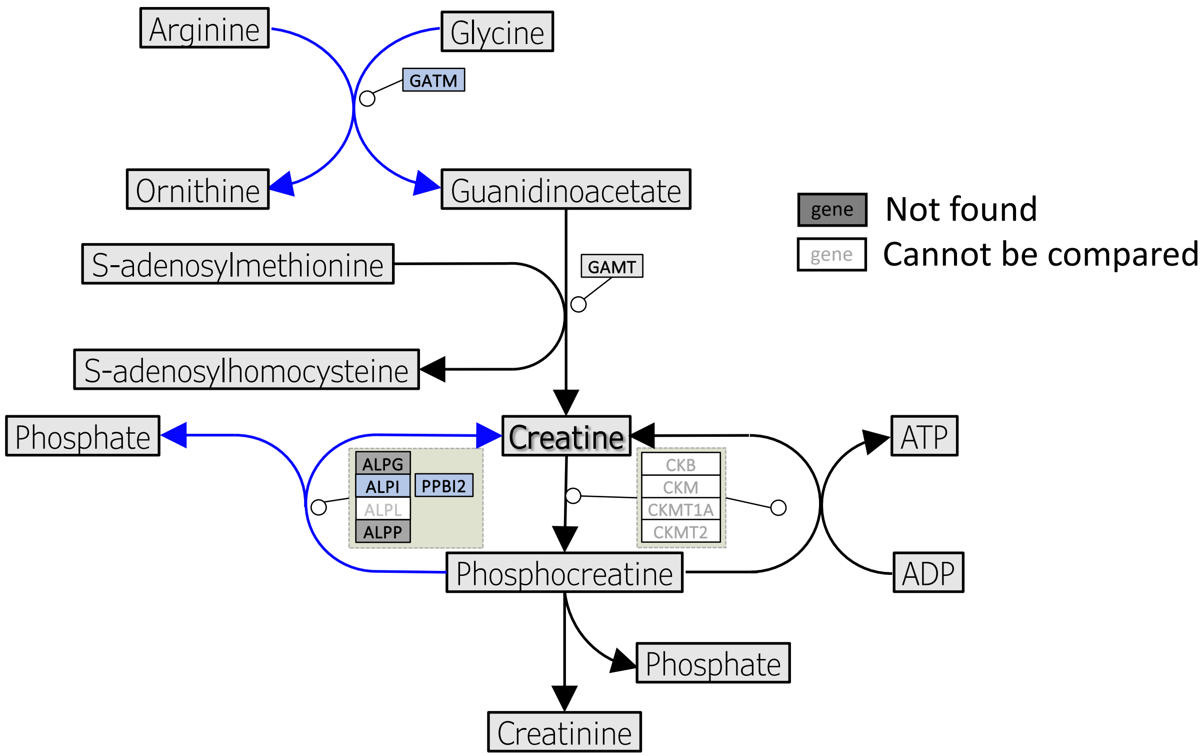

Arginine
Glycine
GATM
Ornithine
Guanidinoacetate
S-adenosylmethionine
GAMT
S-adenosylhomocysteine
Phosphate
Creatine
ATP
ALPG
ALPI
PPBI2
ALPL
ALPP
CKB
CKM
CKMT1A
CKMT2
Phosphocreatine
ADP
Phosphate
Creatinine
gene
Not found
gene
Cannot be compared

### Figure S2

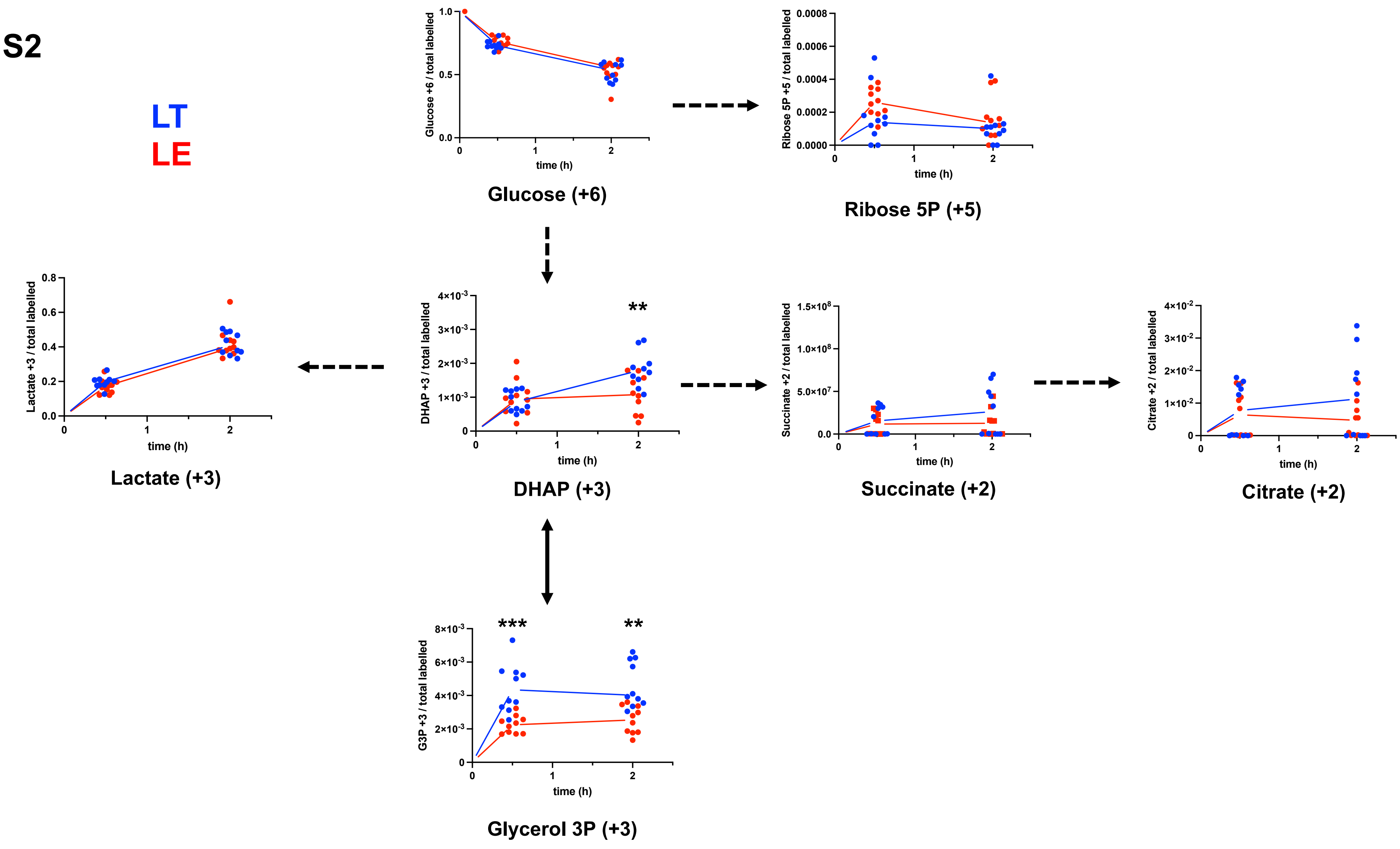
LT
LE
Glucose +6 / total labelled
time (h)
Glucose (+6)
Ribose 5P +5 / total labelled
Ribose 5P (+5)
Lactate +3 / total labelled
Lactate (+3)
DHAP +3 / total labelled
**
DHAP (+3)
Succinate +2 / total labelled
Succinate (+2)
Citrate +2 / total labelled
Citrate (+2)
G3P +3 / total labelled
***
**
Glycerol 3P (+3)

### Figure S3

A

```
1 ATGGCATTTCAGAAGGCAGTTAAAGGGACTATTCTAGTTGGAGGAGGGGCTCTTGCCACAGTTCTAGGACTTTCTCAGTTTGCTCAATAC
1 M A F Q K A V K G T I L V G G G A L A T V L G L S Q F A Q Y

91 AGACGGAAGCAAGTGAACCTGGCGTATGTAGAAGCAGCCGAGTGCTTTACAGAACCTGTCAACCGGGAGCCTCCTTCCAGAGAAGCTCAG
31 R R K Q V N L A Y V E A A E C F T E P V N R E P P S R E A Q

181 CTACTGACTCTGCAAAACACACCTGAATTTGATATCCTTGTTGTTGGAGGAGGAGCAACAGGCAGTGGCTGTGCGCTAGATGCTGTTACT
061 L L T L Q N T P E F D I L V V G G G A T G S G C A L D A V T

271 CGAGGACTAAAAACAGCCCTTGTTGAAAGAGATGATTTCTCATCAGGGACCAGCAGCAGAAGCACTAAATTGATCCATGGTGGTGTGAGA
091 R G L K T A L V E R D D F S S G T S S R S T K L I H G G V R

361 TATCTTCAGAAGGCCATCATGAAGTTGGATATTGAGCAGTATAGGATGGTAAAAGAAGCCCTTTATGAGCGTGCCAACCTACTAGAAATT
121 Y L Q K A I M K L D I E Q Y R M V K E A L Y E R A N L L E I

451 GCTCCCCATTTATCAGCTCCATTACCTATAATGCTTCCAATTTACAAGTGGTGGCAGTTACCTTACTACTGGGTAGGAATCAAGCTATAT
451 GCTCCCCACTTATCAGCTCCATTACCTATAATGCTTCCAATTTACAAGTGGTGGCAGTTACCTTACTACTGGGTAGGAATCAAGCTATAT
151 A P H L S A P L P I M L P I Y K W W Q L P Y Y W V G I K L Y

541 GATCTGGTTGCAGGAAGTAGTTGCTTGAAAAGCAGTTATGTCCTCAGCAAATCGAGAGCCCTTGAACATTTCCCAATGCTCCAGAAAGAC
541 GATCTGGTTGCAGGAAGTAATTGCTTGAAAAGCAGTTATGTCCTCAGCAAATCGAGAGCCCTTGAACATTTCCCAATGCTCCAGAAAGAC
181 D L V A G S S C L K S S Y V L S K S R A L E H F P M L Q K D
181 D L V A G S N C L K S S Y V L S K S R A L E H F P M L Q K D

631 AAATTAGTAGGAGCAATTGTCTACTATGACGGACAACACAACGATGCGCGGATGAACCTTGCCATTGCACTCACTGCTGCCAGGTATGGG
211 K L V G A I V Y Y D G Q H N D A R M N L A I A L T A A R Y G

721 GCTGCCACAGCCAATTACATGGAGGTAGTGAGCTTGCTCAAGAAGACAGACCCTGAGACAGGCAAAGAGCGTGTGTGCGGTGCCCGGTGC
241 A A T A N Y M E V V S L L K K T D P E T G K E R V C G A R C

811 AAGGATGTCCTCACAGGGCAGGAATTTGACGTGAGAGCCAAATGTGTTATCAACGCTACAGGACCTTTCACAGACTCTGTGCGCAAAATG
271 K D V L T G Q E F D V R A K C V I N A T G P F T D S V R K M

901 GATGATAAGAACGCTGCAGCCATCTGCCAGCCGAGTGCTGGTGTCCATATTGTGATGCCTGGTTATTACAGCCCCGAGAGCATGGGACTT
301 D D K N A A A I C Q P S A G V H I V M P G Y Y S P E S M G L

991 CTTGACCCAGCAACCAGTGATGGGCGAGTTATTTTCTTCTTACCCTGGCAAAAGATGACTATTGCTGGCACAACCGATACTCCAACTGAT
331 L D P A T S D G R V I F F L P W Q K M T I A G T T D T P T D

1081 ATCACATCCCATCCTATTCCTTCGGAAGAAGATATTAACTTCATTTTGAATGAAGTGCGTAACTACCTGAGTTGTGATGTGGAAGTGAGA
0361 I T S H P I P S E E D I N F I L N E V R N Y L S C D V E V R

1171 AGAGGAGATGTCCTAGCAGCATGGAGTGGTATCCGTCCCCTCGTTACTGATCCCAACTCTGCAGATACTCAGTCCATCTCTAGAAATCAC
0391 R G D V L A A W S G I R P L V T D P N S A D T Q S I S R N H

1261 GTTGTTGATATCAGTGAGAGTGGCCTTATCACTATAGCAGGTGGAAAATGGACAACTTACCGATCTATGGCGGAAGATACTGTAAATGCT
1261 ATTGTTGATATCAGTGAGAGTGGCCTTATCACTATAGCAGGTGGAAAATGGACAACTTACCGATCTATGGCGGAAGATACTGTAAATGCT
0421 V V D I S E S G L I T I A G G K W T T Y R S M A E D T V N A
0421 I V D I S E S G L I T I A G G K W T T Y R S M A E D T V N A

1351 GCCATCAAGACTCACAATTTGAAAGCAGGACCAAGTAGAACAGTTGGGCTTTTCCTTCAAGGGGGCAAAGATTGGAGCCCCACCCTCTAC
1351 GCCATCAAGACTCACAATTTGAAAGCAGGACCAAGTAGAACAGTTGGGCTTTTCCTTCAAGGAGGCAAAGATTGGAGCCCCACCCTCTAC
0451 A I K T H N L K A G P S R T V G L F L Q G G K D W S P T L Y

1441 ATCCGGCTTGTCCAGGATTATGGACTTGAAAGTGAGGTGGCACAGCATCTTGCTGCCACCTATGGTGACAAGGCTTTTGAGGTGGCCAAA
0481 I R L V Q D Y G L E S E V A Q H L A A T Y G D K A F E V A K

1531 ATGGCAAGTGTGACTGGAAAGAGATGGCCTGTCGTTGGAGTACGTCTTGTGTCAGAATTTCCATATATTGAGGCAGAGGTAAAATACGGT
0511 M A S V T G K R W P V V G V R L V S E F P Y I E A E V K Y G

1621 ATTAAGGAGTACGCCTGCACCGCGGTGGATATGATTTCACGCCGCACTCGTCTGGCCTTTCTCAACGTTCAGGCAGCAGAAGAAGCCCTA
1621 ATTAAGGAGTACGCCTGCACCGCAGTGGATATGATTTCACGCCGCACTCGTCTGGCCTTTCTCAACGTTCAGGCAGCAGAAGAAGCCCTA
0541 I K E Y A C T A V D M I S R R T R L A F L N V Q A A E E A L

1711 CCGAGGATTGTTGAACTGATGGGCAGAGAATTGAATTGGGATGATTATAAGAAAGAGCAAGAACTTGAAACAGCCAGGAAGTTTCTTTAT
0571 P R I V E L M G R E L N W D D Y K K E Q E L E T A R K F L Y

1801 TACGAAATGGGCTATAAATCTCGATCAGAACAATTAACAGATCGCTCTGAAATCAGCCTGCTGCCTTCAGACATCGACAGGTACAAGAAG
0601 Y E M G Y K S R S E Q L T D R S E I S L L P S D I D R Y K K

1891 AGATTTCATAAGTTTGATGCAGACCAAAAAGGATTTATTACCATTGTTGATGTTCAGCGTGTATTAGAGAGTATCAATGTCCAAATGGAT
```

B

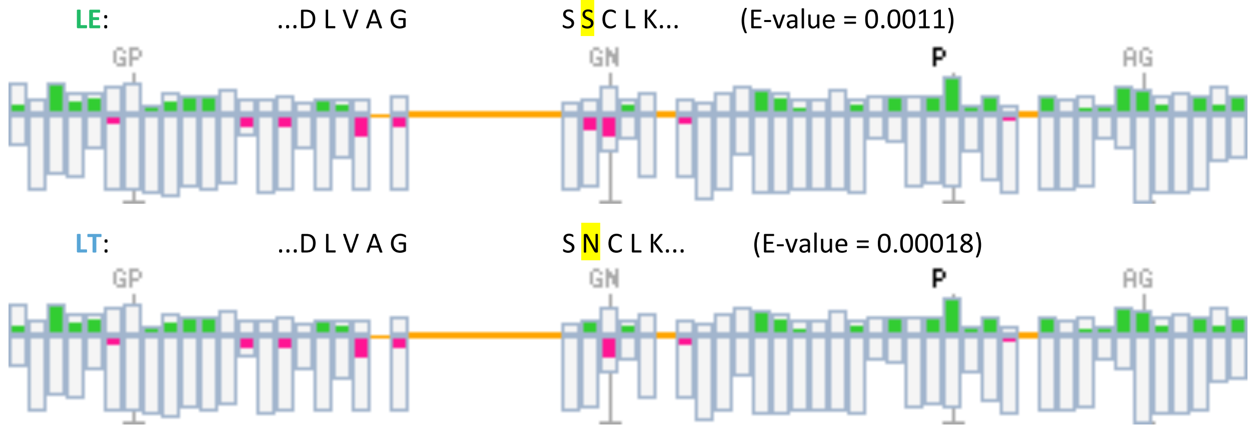

**Figure S4**

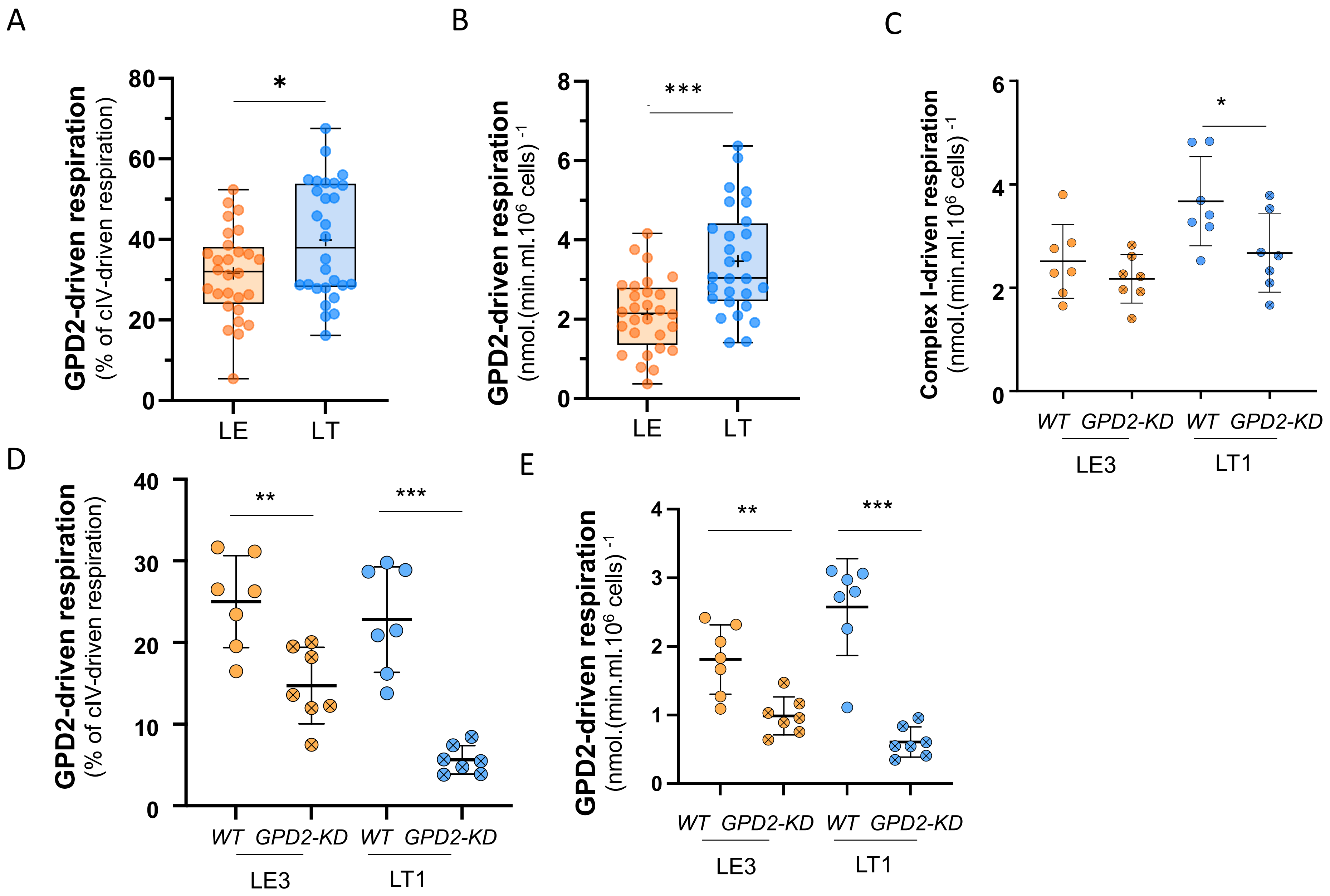
A
GPD2-driven respiration
(% of cIV-driven respiration)
80
60
40
20
0
*
LE
LT
B
GPD2-driven respiration
(nmol.(min.ml.10^6 cells) -1
8
6
4
2
0
***
LE
LT
C
Complex I-driven respiration
(nmol.(min.ml.10^6 cells) -1
6
4
2
0
*
WT GPD2-KD WT GPD2-KD
LE3
LT1
D
GPD2-driven respiration
(% of cIV-driven respiration)
40
30
20
10
0
**
***
WT GPD2-KD WT GPD2-KD
LE3
LT1
E
GPD2-driven respiration
(nmol.(min.ml.10^6 cells) -1
4
3
2
1
0
**
***
WT GPD2-KD WT GPD2-KD
LE3
LT1
